# A physics-informed neural network framework for inferring infinitesimal phase response curves from finite-pulse circadian recordings

**DOI:** 10.64898/2026.10.20.760317

**Authors:** Dongwoo Won, Gyuyoung Hwang, Jae Kyoung Kim

## Abstract

Biological clocks reset and entrain to external cues in a phase-dependent manner. The infinitesimal phase response curve (iPRC) provides the local dynamical rule underlying this process by quantifying how an input changes the instantaneous phase velocity of an oscillator. Inferring iPRCs would make it possible to predict how rhythms respond to new stimulus timings and durations, but existing approaches often require detailed mechanistic models, long perturbation recordings, or specialized population-level resetting experiments. Here we introduce Physics-informed neural network for iPRC inference (PINN-iPRC), a physics-informed neural network framework for inferring iPRCs from standard finite-pulse stimulus-response recordings. PINN-iPRC represents the unknown iPRC as a neural network and embeds the phase equation into the loss function, identifying the iPRC that leads to the observed transient phase trajectory under the known stimulus. In synthetic oscillators, PINN-iPRC accurately recovered iPRCs from finite-pulse recordings, including complex ones that are difficult to capture with simpler parametric reconstructions. Applied to existing experimental Arabidopsis circadian recordings, PINN-iPRC recovered iPRCs comparable to those obtained from specialized population-level resetting experiments, while requiring only conventional stimulus-response trajectories. The inferred iPRCs were further used to predict finite-duration PRCs and circadian peak phases across photoperiods. In particular, the predicted near-invariance of the *prr7* peak phase across 8–16 h of darkness was consistent with independently published *prr7-11* root recordings. These results support PINN-iPRC as a practical physics-informed approach for extracting interpretable phase-sensitivity functions from routine biological oscillator recordings, with potential applications in experimental design, entrainment prediction, and chronotherapeutic or pharmacological schedule optimization.

## Introduction

Biological oscillators are widespread in living systems and coordinate diverse physiological processes, including circadian clocks [1,2], cell cycles [3–5], cardiac rhythms [6–8], and neuronal oscillators [9–11] (Fig. 1(a)). These oscillators are continually shaped by external stimuli such as light [12,13], temperature [14,15], nutrients [16], and biochemical signals [17] (Fig. 1(b)). Because these inputs can reset, stabilize, or synchronize oscillatory dynamics, understanding stimulus-induced phase responses is essential for characterizing how biological rhythms adapt to their environment. A defining feature of these responses is that they depend on the oscillator state at the time of stimulation [18,19]: a circadian clock in the early night is dynamically different from the same clock in the late night, and cardiac or neuronal oscillators can respond differently depending on their position within the cycle. As a result, the same external stimulus can produce qualitatively different effects depending on when it is delivered. In the human circadian clock, for example, light exposure during the early biological night delays the clock, whereas light exposure later in the biological night advances it, and light delivered during insensitive phases produces little phase shift [20].

**Figure 1.**
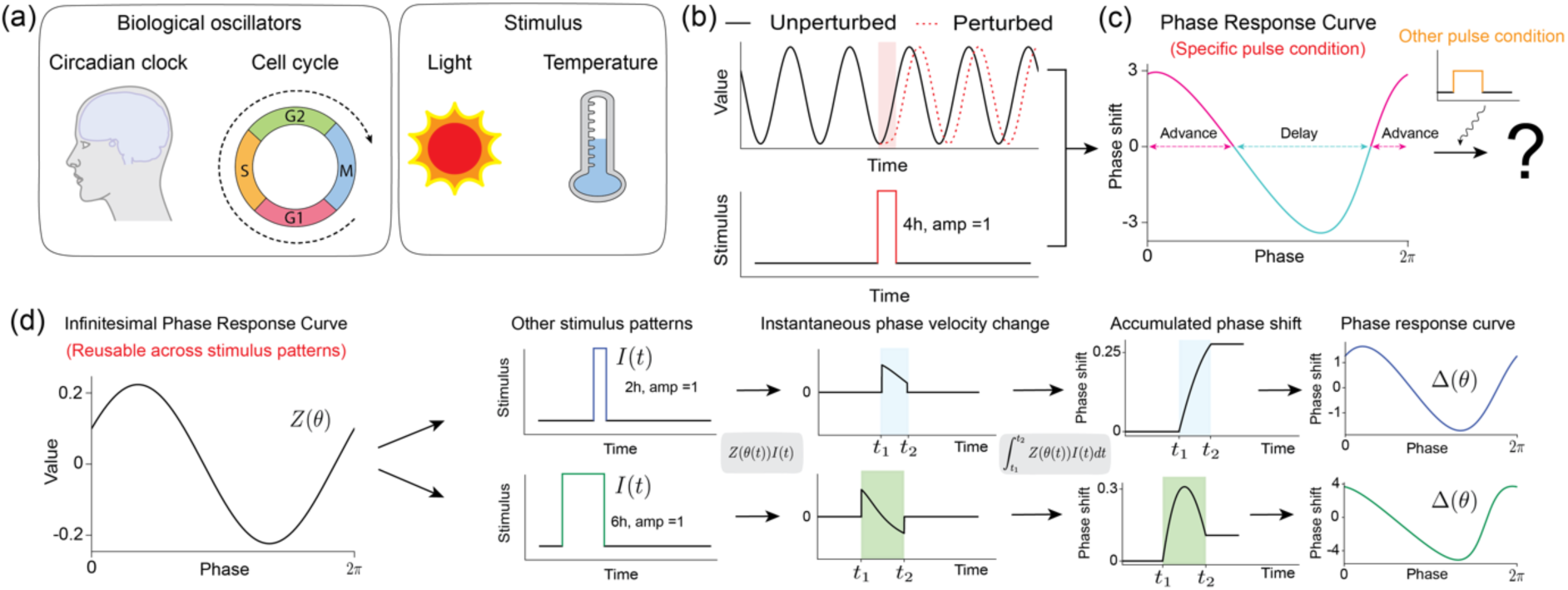
Phase responses of biological oscillators to external stimuli can be characterized by the PRC and iPRC. (a) Biological oscillators are widespread in living systems and appear across diverse biological contexts, including circadian clocks and cell cycles. These oscillators are often exposed to external stimuli such as light and temperature, and understanding how their phase changes in response to such stimuli is essential for characterizing oscillator dynamics. (b) The response of an oscillator to a stimulus can be observed as a time-series trajectory. When a stimulus is applied over a specific time interval, the perturbed oscillation (dotted red line) subsequently proceeds with a shifted phase relative to the unperturbed trajectory. (c) The phase response curve (PRC) summarizes this stimulus-induced final phase shift as a function of the phase at which the stimulus is applied. For a given stimulus condition, a positive PRC value indicates a phase advance, whereas a negative PRC value indicates a phase delay. However, this PRC curve is obtained from a specific pulse condition, and it does not itself provide a local dynamical rule for predicting responses to other pulse conditions. (d) The infinitesimal phase response curve (iPRC) *Z*(*θ*) describes the instantaneous phase sensitivity of an oscillator to a weak perturbation at each phase. When multiplied by the external stimulus function *I*(*t*), the iPRC determines the stimulus-induced change in instantaneous phase velocity. This velocity change is accumulated to yield the total phase shift, thereby producing the phase response curve Δ(θ) corresponding to the stimulus.

This phase dependence is not merely a descriptive feature of oscillatory systems; it has direct physiological and clinical consequences. Patients with bipolar I disorder have been reported to show heightened melatonin suppression in response to nocturnal light compared with healthy individuals, indicating altered circadian light sensitivity [21]. Moreover, light and melatonin interventions must be timed appropriately because their phase-shifting effects depend on when they are administered during the circadian cycle [20,22,23]. Accurate light-response information is also central to model-based circadian interventions, including optimized light schedules for recovery from jet lag [24], shift-work adaptation, and personalized sleep-timing strategies for improving circadian alignment, sleep, or alertness [25–27]. Thus, understanding not only whether an oscillator responds to a stimulus, but also how the direction and magnitude of that response depend on the phase of stimulation, is essential for predicting circadian resetting, optimizing intervention timing, and ultimately characterizing entrainment.

These phase-dependent stimulus responses are commonly quantified by the phase response curve (PRC) [18,28–30] (Fig. 1(c)). A PRC measures the final phase advance or delay caused by a stimulus delivered at different phases, thereby providing a compact description of when the oscillator is delayed, advanced, or minimally shifted by that stimulus [31]. Experimentally, PRCs are typically obtained by applying finite perturbations at multiple phases and measuring the resulting asymptotic phase shifts, although several methods have also inferred PRCs from reconstructed oscillatory trajectories or physiological recordings [32,33]. However, this compact description remains an endpoint summary: it reports the net phase shift after a stimulus has acted, but it does not specify how the phase velocity changed during stimulation. The same final phase shift can arise from different time-resolved phase trajectories. Importantly, a PRC measured for one stimulus amplitude, duration, and temporal profile does not by itself provide a general dynamical rule for predicting responses to other input patterns (Fig. 1(c)). Such prediction requires a local description of how external inputs modify phase velocity at each phase of the oscillator. The infinitesimal phase response curve (iPRC) provides this description by quantifying the instantaneous sensitivity of oscillator phase to external inputs (Fig. 1(d)) [18,19,34,35]. By linking time-varying stimuli to continuous changes in phase velocity within the phase-reduction framework, the iPRC provides a dynamical basis for predicting phase resetting and entrainment across stimulus conditions [36].

Unlike the PRC, however, the iPRC is not directly observed because it describes an instantaneous phase-sensitivity function rather than an endpoint phase shift. The central challenge is therefore to estimate this hidden dynamical quantity from experimentally accessible recordings. Existing approaches can be broadly organized according to the type of information they require about the oscillator and its response to external stimuli (Fig. 2). When a mechanistic state-space model is available, the iPRC can be computed from the governing equations. These equations define a stable limit cycle, along which the oscillator dynamics can be linearized, and the resulting adjoint equation yields the iPRC (Fig. 2(a)) [30,34,36]. This adjoint-based approach is mathematically rigorous because it derives phase sensitivity directly from the vector field governing oscillator dynamics. In real biological systems, however, such mechanistic information is rarely available with sufficient accuracy. Moreover, adjoint-based computation requires access to the full state-space dynamics of the oscillator, whereas experiments often provide only one or a few observable time series, such as reporter signals in circadian assays.

**Figure 2.**
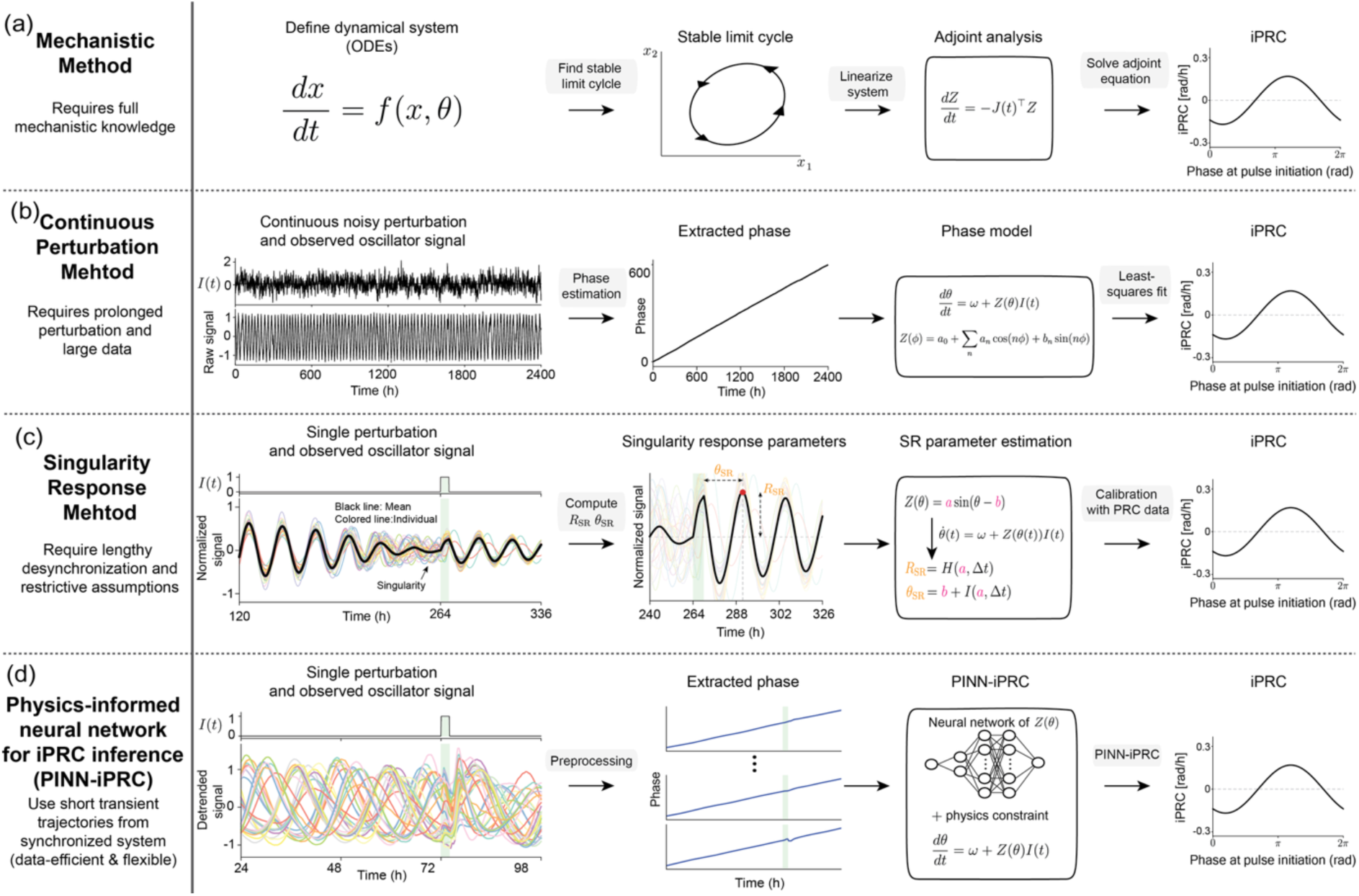
Comparison of iPRC inference methods and the PINN-iPRC framework. (a) Mechanistic method. A dynamical system is specified as a set of ordinary differential equations. A stable limit cycle is obtained from the model, the system is linearized along the limit cycle, and the adjoint equation is solved to compute the iPRC. (b) Continuous perturbation method. A continuously perturbed oscillator signal and the corresponding input *I*(*t*) are observed over time. The oscillatory signal is converted into a phase trajectory, and the iPRC is estimated by fitting a phase model to the extracted phase and the known input. (c) Singularity response method. A single stimulus is applied to a desynchronized oscillator population. The population rhythm after stimulation is summarized by singularity response parameters, including reset phase *θ*_SR_ and reset amplitude *R*_SR_ . These parameters are used together with phase-shift data to estimate the iPRC through an SR parameter estimation procedure. (d) Physics-informed neural network for iPRC inference (PINN-iPRC). Finite-pulse stimuli are applied to synchronized oscillator signals at different initial phases. The observed oscillatory signals are preprocessed to obtain phase trajectories. The extracted phase trajectories and the known stimulus profile are used in the PINN-iPRC model, where the iPRC is represented as a neural network constrained by the phase equation.

When such mechanistic information is unavailable, data-driven trajectory-based approaches can be used. In continuous-perturbation methods, an oscillator is recorded under a time-varying input, the oscillatory signal is converted into a phase trajectory, and the iPRC is estimated by fitting a phase equation to the extracted phase and the known input profile (Fig. 2(b)) [35,37]. Related deep-learning approaches similarly avoid explicit governing equations by learning surrogate oscillator dynamics or latent phase representations from data, from which response functions or phase sensitivity functions can be estimated [38]. However, replacing mechanistic knowledge with data places a greater burden on the recordings themselves: reliable inference requires sufficiently long and informative trajectories, input variability, broad phase coverage, and accurate phase or state reconstruction over time. These requirements can be difficult to satisfy in biological experiments, where perturbations are often delivered as finite pulses and only short stimulus-response trajectories are available.

To reduce the need for long continuously perturbed recordings, singularity response (SR)-based methods use finite-pulse stimulation of desynchronized oscillator populations (Fig. 2(c)) [39–42]. In these methods, a stimulus is applied near a population-level singularity state, and the resulting reset phase and reset amplitude are combined with phase-shift data to estimate the iPRC through a low-dimensional phase model. Although this strategy has been successful in plant and mammalian circadian clocks [39,40], it introduces a different experimental burden: the oscillator population must first be brought to an appropriate desynchronized state, which can require several days in circadian assays, and the stimulus must be delivered at the appropriate population state. In addition, because SR measurements provide only low-dimensional summary quantities, iPRC reconstruction often relies on restrictive assumptions about the phase sensitivity function, such as a sinusoidal form. These requirements can limit the recovery of asymmetric, multi-peaked, or otherwise non-sinusoidal iPRC structures. Although higher-harmonic SR theory can recover more detailed PRC shapes, it requires the individual oscillator waveform to be known and the oscillator population to be near-uniformly desynchronized at the time of stimulation [42].

These limitations raise a natural question of whether iPRCs can be reliably inferred from the relatively short finite-pulse trajectories commonly available in biological experiments. We address this question with Physics-informed neural network for iPRC inference (PINN-iPRC), a physics-informed neural network framework for inferring iPRCs from phase trajectories extracted during and after finite stimulation (Fig. 2(d)). Instead of requiring a mechanistic state-space model or full access to the oscillator’s vector field, PINN-iPRC represents the iPRC as a neural network and constrains its inference with the phase equation using the observed transient phase dynamics and the known stimulus profile. Because this information is extracted from finite-pulse stimulus-response trajectories, PINN-iPRC extracts phase-sensitivity information directly from standard stimulus-perturbed oscillatory recordings. Thus, PINN-iPRC requires neither long continuously perturbed recordings nor desynchronized population-level resetting experiments (SR measurements).

Using this framework, we first tested whether transient phase trajectories contain sufficient information to recover known iPRCs. In synthetic oscillators with ground-truth phase sensitivity functions, PINN-iPRC inferred iPRCs that closely approximated sinusoidal and non-sinusoidal ground-truth functions from finite-pulse stimulus-response trajectories. We then re-analyzed existing Arabidopsis circadian clock recordings from [39], which had originally been analyzed using SR-based reconstruction. Whereas the SR-based approach required specialized SR measurements in addition to conventional phase-shift data, PINN-iPRC used only standard finite-pulse stimulus-response trajectories. Despite relying on this more readily obtainable data type, PINN-iPRC recovered iPRCs similar to those inferred by the SR-based approach. The inferred mutant-specific iPRCs were subsequently used to generate model-based predictions of finite-duration PRCs for 8-, 12-, and 16-h dark intervals and circadian peak phases under the corresponding 24-h light–dark schedules. Notably, the predicted near-invariance of the *prr7* peak phase across these photoperiods was consistent with independent published *prr7-11* root CCA1::LUC recordings [43]. Together, these results show that PINN-iPRC broadens the practical scope of iPRC inference by leveraging a physics-informed neural network to infer dynamical phase-sensitivity information from standard finite-pulse stimulus-response recordings.

## Results

### Design of PINN-iPRC for flexible iPRC inference from finite-pulse transient phase trajectories

To infer iPRCs from finite-pulse stimulus-response recordings, we developed PINN-iPRC, a physics-informed neural network framework that connects the observed phase trajectory, the known stimulus profile, and the unknown iPRC through the phase equation (Fig. 3(a)). This design allows PINN-iPRC to use transient phase dynamics generated during stimulation, rather than relying only on final phase shifts, as dynamical information for iPRC inference (see Methods for details).

**Figure 3.**
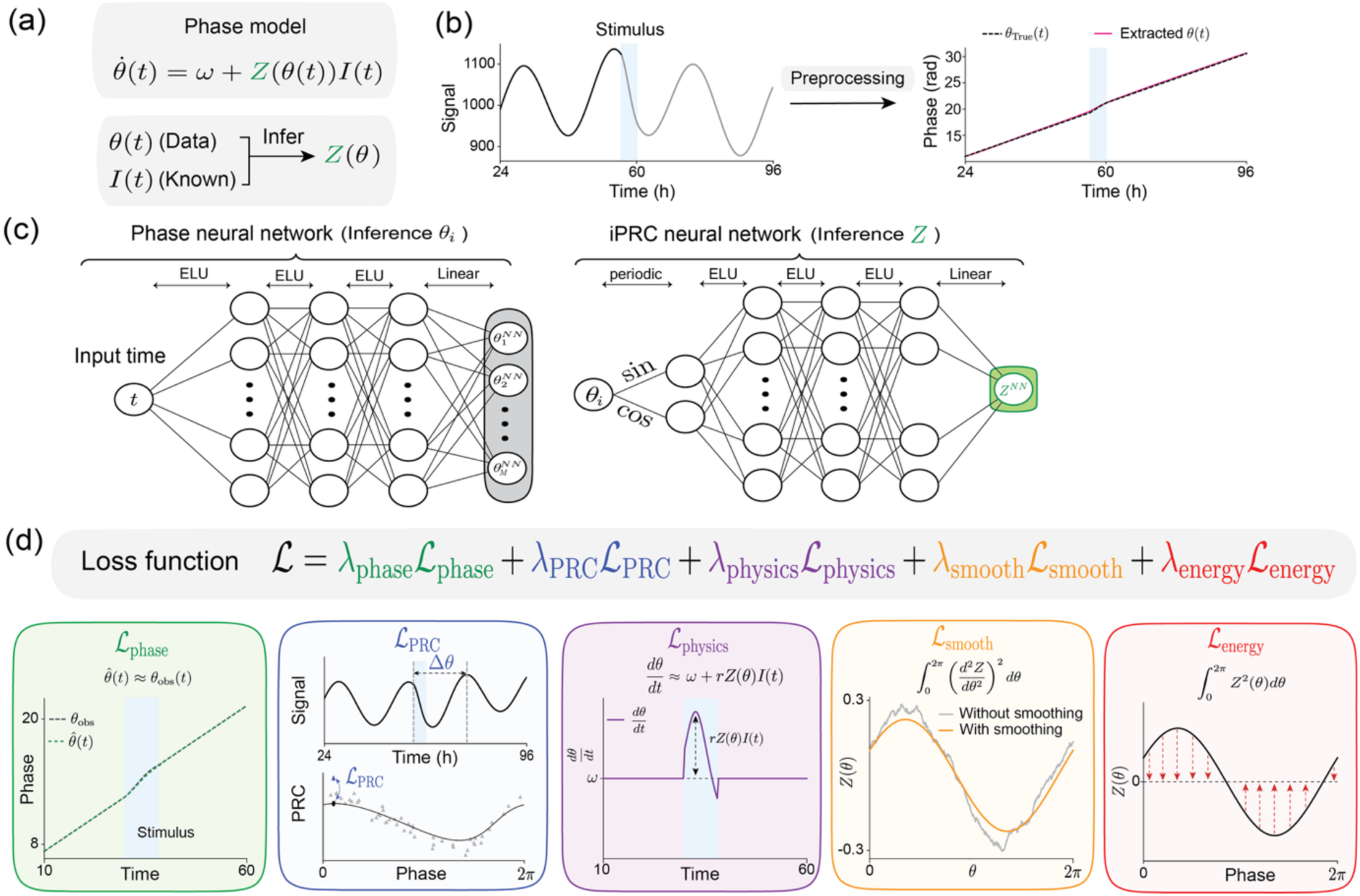
PINN-iPRC framework for inferring the iPRC from stimulus-perturbed oscillatory signals. (a) The phase dynamics of an oscillator with intrinsic angular frequency *ω* under an external stimulus *I*(*t*) are described by a phase equation, in which the iPRC *Z*(*θ*) determines the instantaneous phase sensitivity to the stimulus. PINN-iPRC aims to infer *Z*(*θ*) from the phase trajectory *θ*(*t*) and the given stimulus *I*(*t*). (b) An oscillatory signal changes its trajectory when a stimulus is applied during a finite time interval, indicated by the light-blue shaded region. Through a preprocessing procedure (see Methods for details), the observed signal is transformed into a phase trajectory *θ*(*t*) used for model training. The right panel shows that the preprocessing procedure provides an approximate phase trajectory *θ*(*t*) (pink line) that tracks the true phase trajectory *θ*_True_(*t*) (black dotted line) in this synthetic example. (c) The PINN-iPRC framework consists of two neural networks. The first neural network estimates the phase trajectory *θ_i_*(*t*) for each observed signal as a function of the input time *t*. The number of outputs in this network is set to match the number of observed signals. The second neural network models the iPRC function *Z*(*θ*). To encode the periodic nature of phase, the input phase *θ* is first mapped to (sin *θ*, cos *θ*), which is then passed through hidden layers to produce the estimated iPRC *Z^NN^*(*θ*). In both networks, the exponential linear unit is used as the activation function to promote fast convergence and smooth function estimation. (d) PINN-iPRC estimates the phase trajectories and the iPRC by minimizing a total loss function composed of five terms. Two data losses incorporate complementary information extracted from the same stimulus-response recordings. The phase data loss ℒ_phase_ enforces agreement with the extracted phase trajectories, whereas the PRC data loss ℒ_PRC_ constrains the phase shifts deduced from the inferred iPRC to match endpoint phase-shift estimates derived from the same stimulus-response recordings. The physics loss ℒ_physics_ constrains the estimated phase dynamics through a calibrated physics residual based on θ̇ = *ω* + *rZ*(*θ*)*I*(*t*), where *r* is a trainable calibration factor applied only in the physics residual. The unscaled *Z*(*θ*) is used to compute the reported PRCs and all subsequent predictions. Two additional regularization terms are included for stable inference: a smoothness loss ℒ_smooth_that penalizes large second derivatives of the iPRC, and an energy loss ℒ_energy_ that prevents unrealistically large iPRC amplitudes (see Methods for details).

To obtain the phase information required for this framework, each stimulus-perturbed oscillatory signal was converted into a continuous phase trajectory through a preprocessing procedure (Fig. 3(b)). Because finite-pulse stimulation can drive rapid phase progression while also inducing amplitude modulation and transient waveform distortion, robust phase estimation around the stimulation interval is nontrivial. We therefore designed a trajectory-level preprocessing pipeline to stabilize phase extraction while preserving individual perturbation responses. This pipeline combined linear interpolation onto a finer time grid, detrending to remove slow baseline drift, and phase extraction using the Hilbert transform (see Methods and Supplementary Note 3 for details). The resulting phase trajectories were aligned to stimulus onset and used as temporal training data, while the known stimulus profile defined the external input in the phase equation. Together, these two quantities allow PINN-iPRC to infer phase sensitivity from time-resolved stimulus-response dynamics available in standard finite-pulse recordings (Fig. 3(a)).

Using these preprocessed phase trajectories, PINN-iPRC jointly learns the phase dynamics and the iPRC with two neural networks (Fig. 3(c)). The first neural network represents the phase trajectory of each observed signal as a function of time, whereas the second neural network represents the iPRC as a function of phase. To encode the periodic nature of phase, the phase input is mapped to sine and cosine features before being passed to the iPRC network [44–46]. This architecture allows the inferred iPRC to take flexible shapes, including asymmetric and multi-peaked response profiles, without imposing a predefined functional form. The general strategy of jointly learning oscillator trajectories and an unknown phase-dependent function under a phase-dynamics constraint builds on our previous PINN framework for coupling-function inference [46]. Here, we reformulated this strategy specifically for iPRC inference from finite-pulse perturbation data and introduced additional constraints tailored to this inverse problem.

Because flexible neural representations require appropriate constraints, PINN-iPRC was trained with a composite loss function designed to enforce both agreement with the extracted phase trajectories and consistency with phase dynamics (Fig. 3(d)). The phase data loss fits the estimated phase trajectories to the extracted phases, whereas the PRC data loss constrains the deduced phase shifts using endpoint phase-shift estimates derived from the same stimulus-response recordings. The physics loss requires the estimated phase dynamics to satisfy the phase equation under the known stimulus profile. In addition, smoothness and energy regularization terms suppress rapidly oscillating iPRC estimates and penalize excessively large iPRC amplitudes (see Methods for details). Together, these constraints balance flexibility with dynamical and physiological plausibility, enabling PINN-iPRC to represent complex iPRC structures while reducing spurious solutions driven by noise, phase-extraction uncertainty, or excessive model flexibility.

### Transient phase trajectories support inference of complex synthetic iPRCs

Having established the design of PINN-iPRC, we next tested whether finite-pulse transient phase trajectories contain sufficient information to recover iPRCs with known ground truth. This validation was designed to probe two limitations of endpoint- or parametric reconstruction methods [39,40]. First, methods that impose a low-dimensional sinusoidal phase-sensitivity function may recover simple iPRCs but can fail when the true iPRC is asymmetric, multi-peaked, or otherwise non-sinusoidal. Second, methods that rely primarily on endpoint phase shifts can lose information when stimulation induces phase accumulation beyond one cycle, because the final wrapped phase shift may no longer represent the full transient phase advance. We therefore generated synthetic circadian time series from phase models with three ground-truth iPRCs of increasing difficulty: a sinusoidal iPRC, a multi-peaked iPRC designed to violate the sinusoidal assumption, and an extreme phase-advance case designed to test whether transient unwrapped phase dynamics can resolve responses that endpoint summaries obscure (see Methods for details). In the sinusoidal case, where the ground-truth iPRC matched the type of low-dimensional structure assumed by sinusoidal reconstruction, PINN-iPRC recovered the overall iPRC shape and reproduced the corresponding finite-pulse PRC (Fig. 4(a)). This result shows that the flexible neural representation does not sacrifice accuracy in a simple response regime. We then tested PINN-iPRC on a multi-peaked iPRC, a case designed to violate the sinusoidal assumption. Here, PINN-iPRC recovered the main non-sinusoidal features of the ground-truth iPRC and reproduced the corresponding PRC (Fig. 4(b)), showing that flexibility of the neural representation can support recovery of phase-sensitivity structures that cannot be represented by a predefined sinusoidal form.

**Figure 4.**
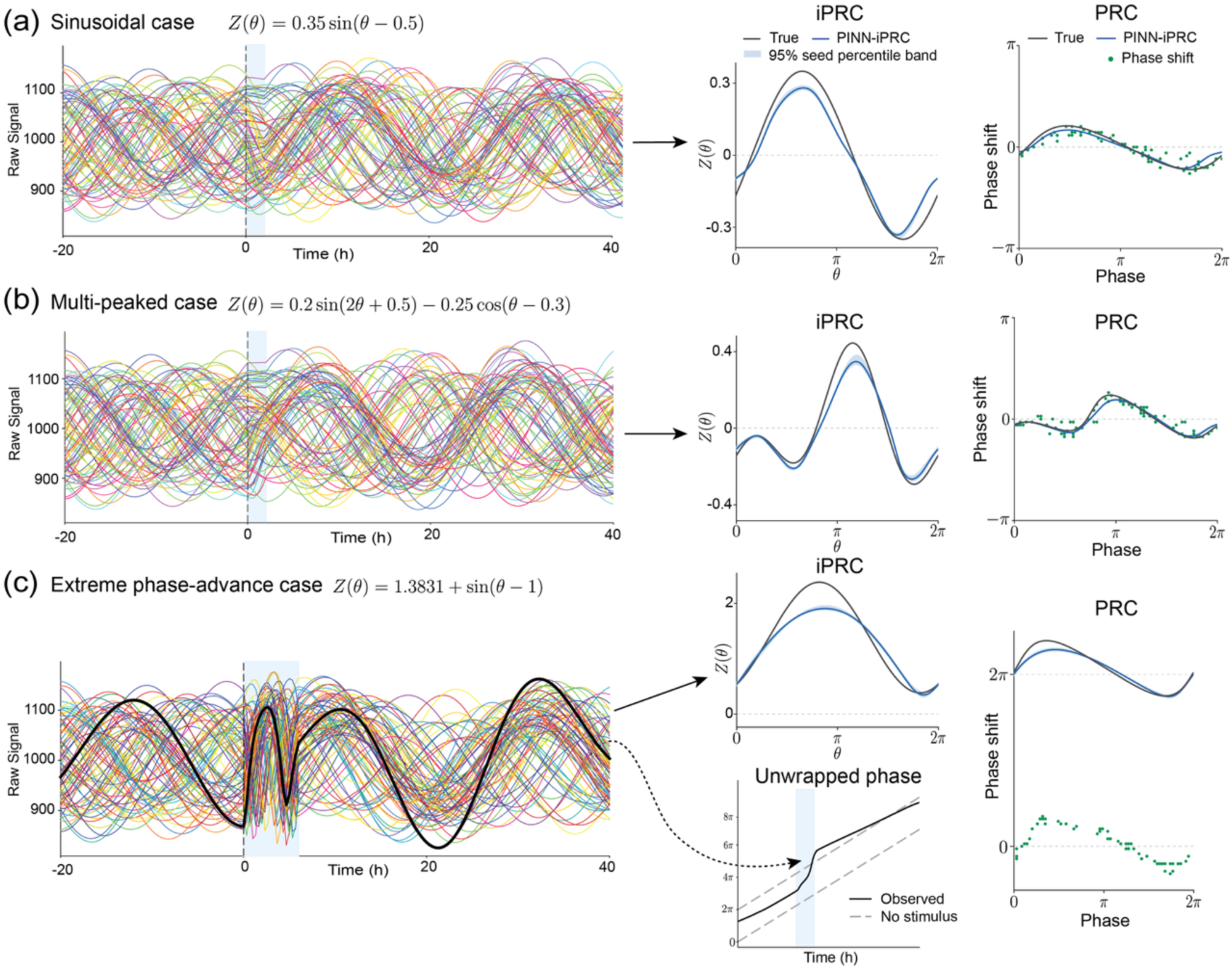
Synthetic benchmarks demonstrate that transient phase trajectories enable recovery of complex iPRCs beyond endpoint-based inference. (a) Synthetic oscillatory signals were generated from a phase model with a sinusoidal ground-truth iPRC and stimulated with a finite pulse at different initial phases (see Methods for details). The left panel shows raw stimulus-response signals. The middle panel compares the ground-truth iPRC (black line) with the PINN-iPRC-inferred iPRC (blue line), which represents the mean estimate across 20 independent training runs, with the shaded region indicating the pointwise 95% seed percentile band (2.5th–97.5th percentiles across the 20 random initializations). The right panel compares the finite-pulse PRC computed from the ground-truth iPRC (black line) with the PRC (blue line) deduced from the mean of PINN-iPRC-inferred iPRCs. Green points indicate phase shifts estimated from the phase difference before and after stimulation. (b) Same as in (a), but using a multi-peaked ground-truth iPRC. (c) Same as in (a), but for an extreme phase-advance regime in which finite-pulse stimulation induced phase accumulation beyond one cycle. An additional panel shows an example of an unwrapped phase trajectory during the stimulus-response interval. In (c), the ground-truth and model-predicted PRCs are unwrapped, whereas the green phase-shift estimates are wrapped to [−*π*, *π*).

Finally, in the extreme phase-advance case, finite stimulation caused rapid phase progression beyond one full cycle, which was visible in the unwrapped transient phase trajectory. Even in this strongly resetting regime, PINN-iPRC captured the dominant iPRC structure and reproduced the corresponding PRC (Fig. 4(c)). This case further illustrates why transient phase trajectories provide information beyond endpoint phase shifts. Endpoint-based estimates summarize only the phase difference before and after stimulation. When stimulation drives phase accumulation beyond one cycle, the final wrapped phase shift can obscure the full unwrapped phase advance and therefore misrepresent the actual finite-pulse response. In contrast, PINN-iPRC uses the time-resolved unwrapped phase trajectory during stimulation, allowing it to recover phase-sensitivity information that would be lost in endpoint-based reconstruction.

To benchmark PINN-iPRC against alternative reconstruction strategies, we compared it with sinusoidal and higher-order Fourier parametric reconstructions, as well as a Tokuda-type Fourier phase-dynamics fitting approach [47] (Supplementary Figs. S3 and S4). Simpler approaches performed well when their underlying assumptions were appropriate, whereas PINN-iPRC showed good recovery across all three synthetic response regimes, including the extreme phase-advance case in which the endpoint-based parametric fits failed after phase accumulation exceeded one full cycle. We further applied the iterative Fourier least-squares method of Cestnik and Rosenblum [35] to the same finite-pulse records (Supplementary Fig. S8). When sufficiently long records were available, both methods accurately recovered the ground-truth iPRC. However, as the number of observed cycles was reduced, the Cestnik– Rosenblum reconstruction deteriorated more rapidly, whereas PINN-iPRC retained lower reconstruction error over shorter records. Thus, the advantage observed here was not a universal improvement in reconstruction accuracy, but greater robustness to reduced observation length within the tested benchmarks.

To examine robustness under experimentally relevant data limitations, we varied the number of stimulated phases, measurement noise, and phase coverage (Supplementary Fig. S7). PINN-iPRC remained stable across the tested sample sizes and measurement-noise levels. Reconstruction error increased when large gaps were introduced into phase coverage, indicating that broad phase coverage remains important for accurate inference.

Despite this overall performance, the inferred iPRCs showed residual deviations from the ground-truth functions. A major source of error was the reconstruction of latent phase from reporter-like oscillatory signals. Rapid stimulus-induced phase progression, amplitude modulation, and transient waveform distortion can introduce errors that remain even after preprocessing. This was already evident in the PRC points estimated directly from the extracted phases, which did not always coincide with the ground-truth PRCs before PINN-iPRC inference (Fig. 4(a)–(c), right panels). Consistently, inference using the true rather than reconstructed phase trajectories substantially reduced both reconstruction error and amplitude attenuation (Supplementary Figs. S1 and S5; Supplementary Table S2). Regularization sensitivity analyses further showed that increasing the smoothness penalty primarily suppressed small-scale fluctuations, whereas sufficiently strong energy regularization reduced the overall iPRC magnitude (Supplementary Fig. S6). Thus, phase-reconstruction error was the primary source of the remaining amplitude attenuation, with excessive regularization providing an additional source. Overall, these benchmarks show that PINN-iPRC can recover the dominant structures of complex iPRCs from finite-pulse transient phase trajectories, while also identifying phase-reconstruction error as the principal practical bottleneck under rapid or strongly resetting responses.

### PINN-iPRC infers iPRCs from real plant circadian recordings without singularity-response measurements

We next assessed whether PINN-iPRC could infer iPRCs from experimental circadian recordings for which ground-truth iPRCs are unavailable. We analyzed Arabidopsis CCA1::LUC recordings from *prr5*, *prr9*, and *prr7* plants exposed to a 2-h dark pulse; these data were previously used for SR-based iPRC reconstruction [39] (see Methods for details). The datasets contain two types of experimental information: conventional PRC measurement trajectories, in which synchronized plants were stimulated at different circadian phases, and SR recordings, in which a desynchronized, low-amplitude oscillator population was stimulated near a singularity state. PINN-iPRC used only the conventional trajectories, whereas the published SR-based reconstruction used conventional phase-shift information together with the additional SR measurements. We therefore treated the SR-based reconstruction as a comparative experimental reference rather than as ground truth.

For each mutant, we trained PINN-iPRC using 20 independent random initializations. Figure 5 shows the pointwise arithmetic mean of the inferred iPRCs across these runs as the central estimate, together with the pointwise 95% seed percentile band (2.5th–97.5th percentiles), which summarizes seed-to-seed optimization variability conditional on the fixed dataset and training protocol. The corresponding deduced PRC was calculated by integrating the phase model under the same 2-h dark-pulse condition using the ensemble-mean iPRC and was compared with the experimentally measured phase shifts.

**Figure 5.**
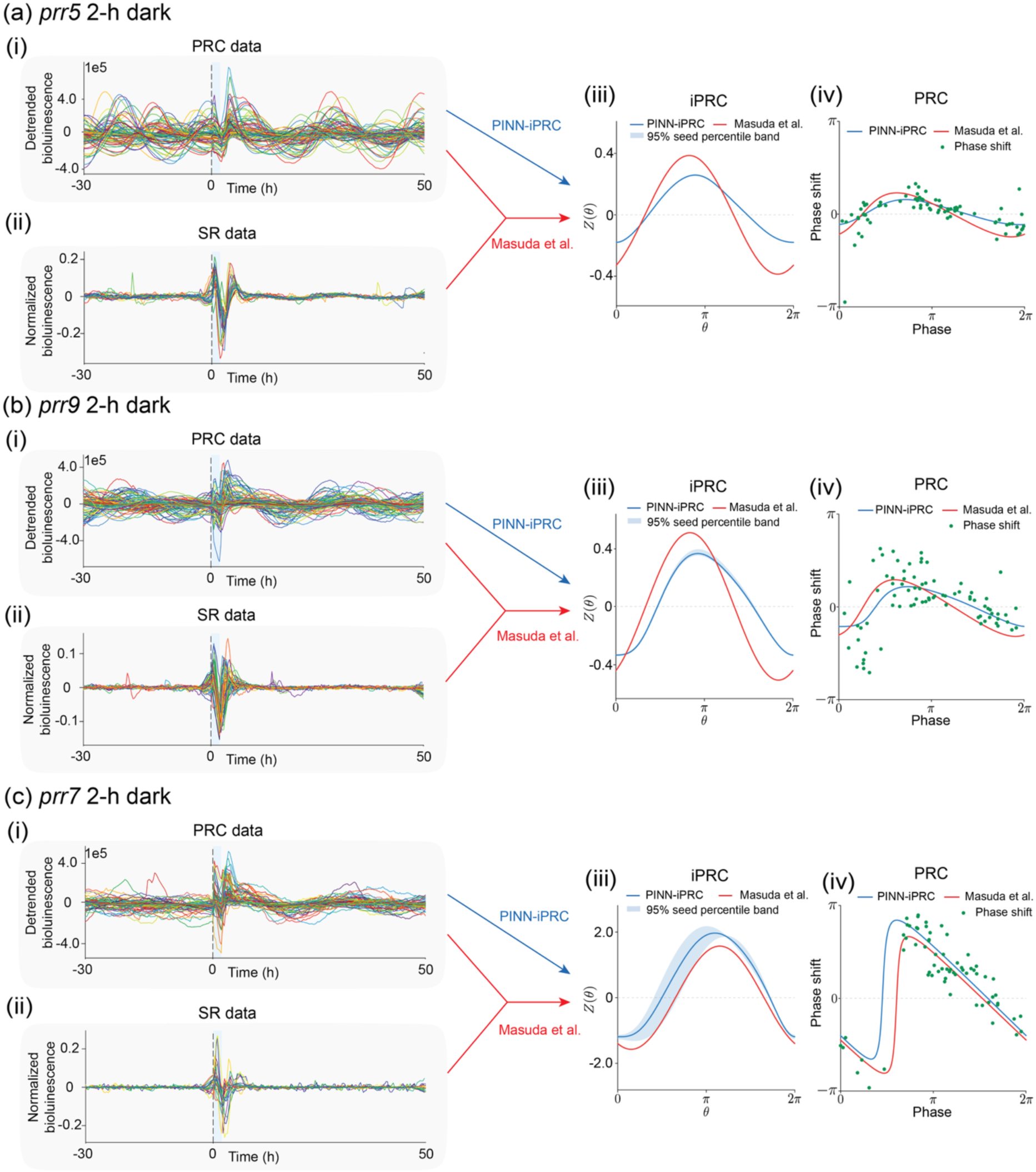
PINN-iPRC infers mutant-specific iPRCs from conventional Arabidopsis dark-pulse trajectories without using singularity-response data. (a) *prr5* CCA1::LUC reporter plants exposed to a 2-h dark pulse. (i) Conventional PRC measurement trajectories are presented, in which a dark pulse was applied to synchronized plants at different circadian phases and the resulting phase shifts were measured from the post-stimulus rhythms. (ii) Singularity-response (SR) trajectories are presented, in which a dark pulse was applied to a desynchronized, low-amplitude oscillator population and the post-stimulus reset response was summarized by SR quantities (Fig. 2(c)). These SR recordings were used for the SR-based reconstruction but not for PINN-iPRC inference. (iii) From the PRC trajectories in (i), the PINN-iPRC-inferred iPRC (blue line) represents the pointwise mean estimate across 20 independent training runs, with the shaded region indicating the pointwise 95% seed percentile band (2.5th– 97.5th percentiles across the 20 random initializations). The SR-based iPRC (red line) was reconstructed using both the conventional PRC measurements in (i) and the SR information in (ii). (iv) The deduced PRCs were obtained by solving the phase model under the 2-h dark-pulse condition using the ensemble-mean iPRC inferred by PINN-iPRC and the published SR-based iPRC, respectively. The phase shifts (green dots) indicate experimentally measured phase shifts obtained from pre- and post-stimulus phase estimates (see Methods for details). (b,c) The same analyses for *prr9* (b) and *prr7* (c) CCA1::LUC reporter plants carrying mutations exposed to a 2-h dark pulse.

In *prr5*, PINN-iPRC inferred a smooth, single-peaked iPRC similar to the SR-based reconstruction, and the deduced PRC reproduced the small Type-1-like phase shifts (Fig. 5(a)). In *prr9*, the inferred iPRC was also similar in overall shape to the SR-based estimate but had a somewhat larger response amplitude than that of *prr5*, resulting in a correspondingly larger deduced PRC (Fig. 5(b)). In *prr7*, which exhibits a large-magnitude Type-0 phase response, PINN-iPRC inferred a substantially larger-amplitude iPRC, consistent in overall shape with the SR-based reconstruction, and the deduced PRC captured the abrupt phase transition characteristic of Type-0 resetting (Fig. 5(c)).

Quantitative comparison further supported agreement in shape and peak phase. Pearson correlations between the ensemble-mean PINN-iPRC and SR-based estimates were 0.956, 0.865, and 0.987, and the corresponding peak-phase differences were 0.226, 0.276, and 0.176 rad for *prr5*, *prr9*, and *prr7*, respectively. The inferred amplitudes were lower than the SR-based estimates for *prr5* and *prr9*, with amplitude ratios of 0.567 and 0.689, respectively, but were comparable for *prr7*, with an amplitude ratio of 1.005 (Supplementary Table S1). Because the seed percentile bands characterize optimization variability rather than uncertainty arising from the finite experimental sample, we separately performed bootstrap resampling of the original trajectories. The resulting bootstrap intervals were narrower for *prr5* and wider for *prr9* and *prr7* (Supplementary Fig. S2).

As a complementary methodological comparison, we applied sinusoidal and higher-order Fourier endpoint-based reconstructions, as well as a Tokuda-type phase-dynamics fitting approach, to the same Arabidopsis datasets (Supplementary Figs. S3 and S4). The sinusoidal reconstructions were broadly similar to the SR-based estimates, whereas the higher-order Fourier and Tokuda-type approaches yielded different iPRC profiles. Because the true experimental iPRCs are unknown, these comparisons characterize agreement and method dependence among the reconstructions but cannot establish their relative accuracy.

Together, these results demonstrate the feasibility of inferring mutant-specific iPRCs from conventional finite-pulse stimulus-response trajectories without requiring additional SR measurements. Although the absence of experimental ground truth precludes ranking the reconstruction methods using these Arabidopsis data, the agreement in overall shape and peak phase with the SR-based reference supports PINN-iPRC as a practical approach when specialized desynchronization and singularity-response experiments are difficult to perform.

## PINN-iPRC predicts mutant-specific responses to prolonged darkness and peak timing across photoperiods

A central advantage of the iPRC is that it provides a local phase-sensitivity function that can be reused with the phase model to compute finite-pulse responses under stimulus conditions that were not directly measured. To examine this predictive capability, we used the PINN-iPRC-inferred iPRCs obtained from the 2-h dark-pulse trajectories of *prr5*, *prr9*, and *prr7* CCA1::LUC plants (Fig. 6(a)). For each mutant, the intrinsic angular frequency was set to *ω* = 2*π*/*τ* using the corresponding free-running period reported by [39] (*τ* = 21.27, 23.42, and 22.85 ℎ for *prr5*, *prr9*, and *prr7*, respectively). For a given initial phase, the finite-duration phase response was computed by integrating the inferred iPRC along the evolving phase trajectory during darkness (Fig. 6(b)). We applied this procedure to dark intervals lasting 8, 12, or 16 h, all of which were longer than the 2-h condition used for iPRC inference, and obtained the corresponding model-predicted PRCs (Fig. 6(c)). These extrapolations assume that darkness is represented by the same effective model input as in the 2-h inference condition and that this input remains unchanged throughout the dark interval, without additional adaptation, desensitization, or history dependence. This assumption does not imply that the resulting phase shift scales linearly with dark duration, because the oscillator continuously traverses different regions of the iPRC during darkness [48,49] (Fig. 6(b)).

**Figure 6.**
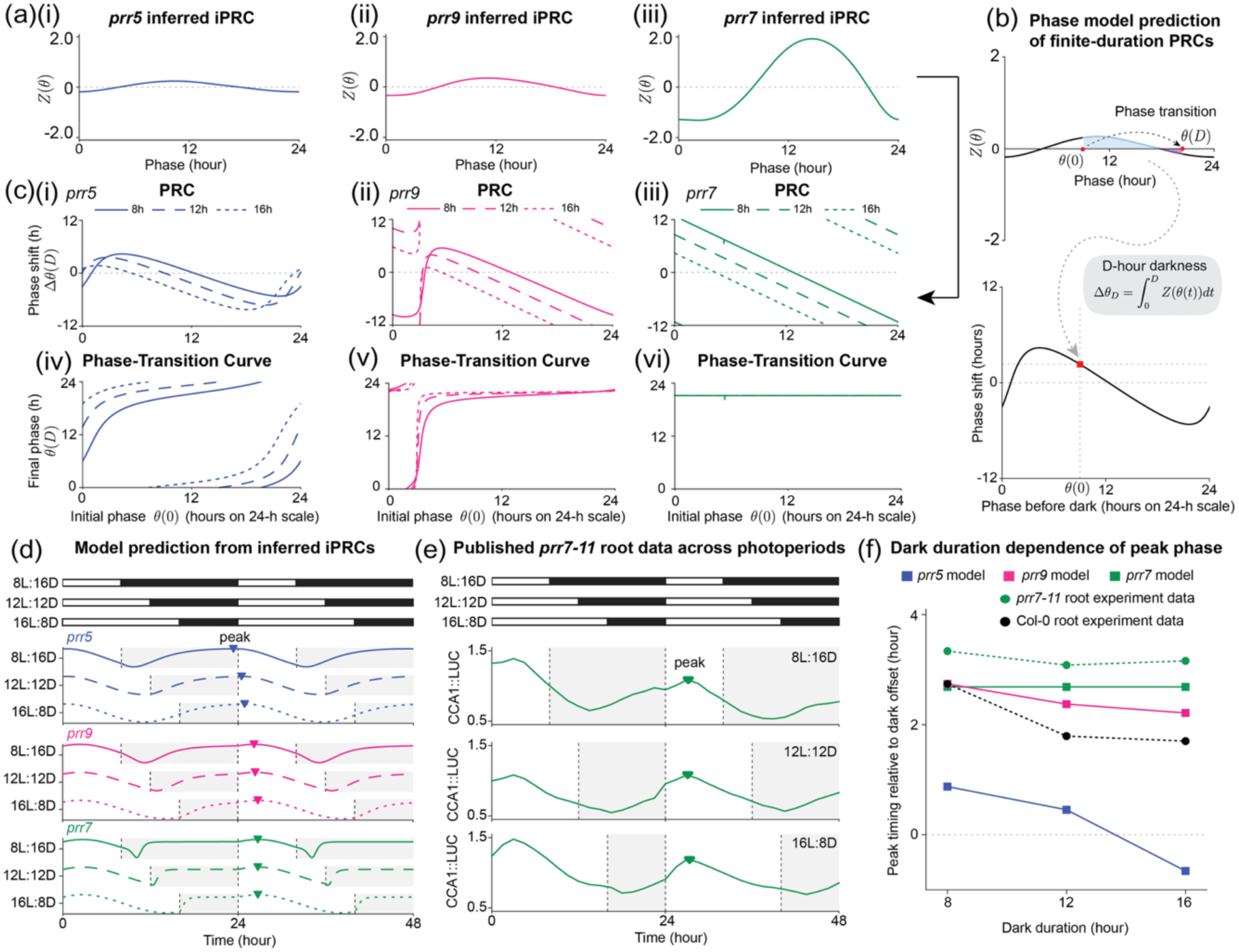
PINN-iPRC-inferred iPRCs generate mutant-specific predictions for prolonged darkness and peak timing across photoperiods. (a) PINN-iPRC-inferred iPRCs obtained from conventional 2-h dark-pulse trajectories of *prr5*, *prr9*, and *prr7* CCA1::LUC plants (Fig. 5) and used in the subsequent simulations. Panels (i)–(iii) show the inferred iPRCs for *prr5*, *prr9*, and *prr7*, respectively. Phase is expressed in hours on a 24-h scale. (b) Schematic of finite-duration PRC calculation from an inferred iPRC. For an oscillator exposed to *D* hours of darkness beginning at phase *θ*(0), the phase evolves according to the phase equation using the inferred iPRC and reaches *θ*(*D*) at the end of the dark interval. The stimulus-induced phase shift relative to unperturbed phase is obtained as Δ*θ_D_* = ∫*^D^* *Z*(*θ*(*t*)>*I*(*t*)*dt*, where *I*(*t*) = 1 during darkness. Repeating this calculation over different initial phases produces the corresponding finite-duration PRC (lower panel). (c) Model-predicted PRCs (i)–(iii) and phase-transition curves (iv)–(vi) for *prr5*, *prr9*, and *prr7*, respectively, under 8-h (solid), 12-h (dashed), or 16-h (dotted) darkness. Phase shifts are expressed in hours and wrapped to the interval [−12,12) on a 24-h scale. Each transition curve maps dark-onset phase to dark-offset phase on a 0–24 h scale. (d) Model-predicted circadian rhythms during two test cycles of 8L:16D, 12L:12D, or 16L:8D following 21 days of 12L:12D preconditioning. White and black bars denote light and dark intervals, respectively. The corresponding mutant-specific iPRC was used in each simulation. Predicted traces are shown for *prr5*, *prr9*, and *prr7* and vertically offset for visualization. Triangles indicate the peak phases used to quantify peak timing relative to dark offset. (e) Digitized published population-mean root CCA1::LUC bioluminescence traces from *prr7-11* plants under the corresponding 8L:16D, 12L:12D, and 16L:8D conditions [43]. These recordings were obtained in an independent experiment and were not used for PINN-iPRC inference. Green triangles indicate the peaks used to estimate peak timing relative to dark offset. (f) Peak phase relative to the first test-dark offset (*t* = 24 ℎ) as a function of dark duration. For both model and experiment, the peak between the first and second dark onsets was selected for each condition; the initial peak was excluded. Negative values indicate peaks before this dark offset. Squares show model predictions obtained from the inferred *prr5*, *prr9*, and *prr7* iPRCs. Green and black circles show peak phases extracted from published *prr7-11* and Columbia-0 (Col-0) wild-type root recordings [43], respectively. Predicted *prr7* peak timing and experimental *prr7-11* peak timing varied only weakly across photoperiods, whereas peak timing in Col-0 varied more substantially.

The predicted PRCs exhibited distinct duration-dependent patterns across the three mutants. In *prr5* and *prr9*, increasing the dark duration altered both the magnitude and shape of the phase response (Fig. 6(c)(i),(ii)). In contrast, *prr7* exhibited a broad phase-shift range and sharp wrapping transitions across all three dark durations, consistent with its strong Type-0-like resetting structure (Fig. 6(c)(iii)). These results illustrate how differences among the inferred mutant-specific iPRCs can produce qualitatively different phase responses when the duration of darkness is extended beyond the inference condition.

We next examined how these duration-dependent responses translated into phase mapping across the dark interval. The corresponding phase-transition curves map the phase at dark onset to that at dark offset (Fig. 6(c)(iv)–(vi)). Unlike *prr5* and *prr9*, the *prr7* curves were nearly horizontal over broad phase ranges, indicating that a wide range of initial phases was mapped to similar post-dark phases. This suggested that *prr7* peak timing would remain similar relative to dark offset across photoperiods.

We then used the inferred iPRCs to simulate circadian rhythms under 8L:16D, 12L:12D, and 16L:8D. For each condition, the simulation consisted of 21 days of 12L:12D preconditioning followed by two cycles of the test photoperiod (Fig. 6(d)). We quantified the peak associated with the first test-dark interval relative to its dark offset at *t* = 24 h. The predicted peak phase of *prr5* changed substantially with dark duration, while *prr9* showed a smaller but still apparent shift. In contrast, the predicted peak phase of *prr7* remained nearly unchanged across the three photoperiods (see Supplementary Note 14 for details of simulation settings and peak-selection criteria).

To assess whether this *prr7* prediction was consistent with independent experimental observations, we analyzed published root CCA1::LUC recordings from *prr7-11* plants obtained under the corresponding 8L:16D, 12L:12D, and 16L:8D conditions [43] (Fig. 6(e)). These data, which were not used for PINN-iPRC inference, similarly showed peaks occurring at nearly the same time relative to dark offset across the three dark durations. Quantification of the peak phases showed that the *prr7* model and the published *prr7-11* root data exhibited both comparable peak timing relative to dark offset and similarly weak dependence on dark duration (Fig. 6(f)). In contrast, the corresponding Col-0 root recordings showed substantially greater peak-phase variation across photoperiods, suggesting that the near-invariant pattern in *prr7-11* was not simply imposed by the photoperiodic schedules themselves.

We further tested whether this model prediction was sensitive to the assumed free-running period. Varying *τ* over the reported *prr7-11* root free-running periods shifted the absolute predicted peak time by up to 0.509 ℎ but did not alter our central prediction that *prr7* peak timing remains nearly invariant across photoperiods (maximum difference < 0.001 h at each fixed *τ*; Supplementary Table S4). Together, these observations provide partial cross-condition support for the predicted weak photoperiod dependence of *prr7* peak timing. However, they do not constitute matched-condition validation of the complete 8-, 12-, and 16-h PRCs. The corresponding predictions for *prr5* and *prr9* remain experimentally untested.

## Discussion

The iPRC provides a local dynamical description of how external inputs modify oscillator phase and is therefore central to predicting resetting and entrainment beyond a single stimulus condition (Fig. 1). However, inferring this local phase-sensitivity function from biological recordings remains difficult because existing approaches typically require a full mechanistic state-space model, long continuously perturbed recordings, or specialized singularity-response measurements (Fig. 2). In this study, we developed PINN-iPRC, a physics-informed neural network framework that infers iPRCs from finite-pulse stimulus-response trajectories (Fig. 3). The key idea of PINN-iPRC is to use the phase equation as a physical constraint: by embedding this equation into the learning objective, PINN-iPRC converts transient phase trajectories into dynamical information about the unknown iPRC. This PINN-based formulation allows a hidden phase-sensitivity function to be inferred from experimentally accessible perturbation recordings without requiring the full oscillator vector field. Across synthetic and experimental circadian datasets, PINN-iPRC recovered flexible iPRCs from transient phase trajectories (Fig. 4) and inferred Arabidopsis mutant-specific iPRCs without requiring singularity-response measurements (Fig. 5). The inferred iPRCs were further used to predict finite-duration PRCs and photoperiod-dependent peak phases, with the predicted *prr7* peak phases showing consistency with independently published *prr7-11* root recordings [43] (Fig. 6). Together, these results support PINN-iPRC as a practical route for extracting interpretable phase-sensitivity functions from routine biological oscillator recordings and using them to generate model-based predictions of resetting and photoperiod-dependent timing beyond the measured stimulus condition.

A central methodological contribution of PINN-iPRC is that it formulates iPRC inference as a physics-informed inverse problem in reduced phase dynamics [46,50,51]. Finite-pulse stimulus-response recordings provide only partial observations of the oscillator: they contain time-resolved trajectories of an observable reporter signal, but not the full molecular state-space dynamics. If the iPRC were learned only by fitting endpoint phase shifts, many different phase-sensitivity functions could produce similar net PRCs, particularly when the data are sparse, noisy, or restricted to a single pulse condition. Conversely, an unconstrained neural network fitted directly to extracted phase trajectories could absorb noise, preprocessing artifacts, or waveform distortions into the inferred response function. PINN-iPRC addresses this ill-posedness by embedding the phase equation into the learning objective, thereby linking the observed transient phase evolution, the known stimulus profile, and the unknown iPRC (Fig. 3). The inferred neural function is therefore not merely a flexible curve fitted to data; it must act as a phase-sensitivity function that explains how the oscillator phase evolves under stimulation. In this sense, PINN-iPRC occupies a middle ground between fully mechanistic adjoint-based inference and unconstrained data-driven fitting: it does not require the full oscillator vector field, but the learned function remains constrained to act as an iPRC within the phase-reduction framework [34,48,49,52,53]. Thus, finite-pulse perturbation recordings, although limited relative to full state-space measurements, can constrain local phase-sensitivity functions when interpreted through a physics-informed phase model.

The synthetic benchmarks allowed us to assess recovery of known phase-sensitivity functions rather than agreement with final phase shifts alone (Fig. 4). Comparisons with endpoint-based reconstructions highlighted the value of both flexible function representation and transient phase information. PINN-iPRC recovered multi-peaked iPRCs beyond a predefined sinusoidal form [39,40]. Higher-order Fourier fitting also captured the multi-peaked structure, but both sinusoidal and higher-order endpoint-based fits failed in the extreme phase-advance case (Supplementary Fig. S3). PINN-iPRC recovered the dominant iPRC structure in this case using transient unwrapped phase information that endpoint phase wrapping can obscure [54].

To distinguish the benefit of the PINN formulation from that of transient phase information itself, we next compared PINN-iPRC with alternative methods that also used trajectory-level information. Tokuda-type Fourier phase-dynamics fitting [47] used the same reconstructed phase trajectories as PINN-iPRC and performed comparably in the extreme case. However, PINN-iPRC showed a smaller amplitude discrepancy in the multi-peaked benchmark (Supplementary Fig. S4), indicating more accurate amplitude recovery for this tested complex iPRC. A comparison with the Cestnik–Rosenblum method [35] using matched oscillatory records further examined robustness to observation length (Supplementary Fig. S8). PINN-iPRC and the Cestnik–Rosenblum method accurately recovered the ground-truth iPRC from sufficiently long records. As the number of observed cycles decreased, however, the Cestnik–Rosenblum reconstruction deteriorated more rapidly, whereas PINN-iPRC retained lower reconstruction error over the shorter records tested here.

The Arabidopsis analyses further showed that the same principle can be applied to real circadian recordings (Fig. 5). Using only conventional finite-pulse stimulus-response trajectories, PINN-iPRC inferred mutant-specific iPRCs that were consistent with those obtained by SR-based reconstruction [39], which requires specialized population-level resetting measurements. A strictly matched-data comparison with higher-harmonic SR methods is less straightforward because these approaches use different experimental observables: higher-harmonic SR reconstruction requires a sufficiently desynchronized population together with information about the individual oscillator waveform [42], whereas PINN-iPRC uses transient phase trajectories from finite-pulse recordings. Accordingly, we regard the SR-based reconstruction here as a comparative experimental reference rather than a matched-data benchmark. Thus, PINN-iPRC broadens the experimental accessibility of iPRC inference by extracting phase-sensitivity information from standard perturbation recordings that would otherwise require stronger modeling assumptions or more specialized experiments.

The mutant-specific analyses illustrate how an inferred iPRC can be used to generate hypotheses beyond the stimulus condition used for inference (Fig. 6). When the iPRCs inferred from 2-h dark-pulse trajectories were applied to longer dark intervals, the three mutants showed distinct duration-dependent responses. In particular, *prr7* was predicted to maintain nearly constant peak timing relative to dark offset across 8L:16D, 12L:12D, and 16L:8D, whereas *prr5* and *prr9* showed greater photoperiod-dependent shifts. This *prr7* pattern was preserved when the assumed free-running period was varied, indicating that it arose from the inferred phase-sensitivity structure rather than from a particular choice of *τ* (Supplementary Table S4). Independent *prr7-11* root CCA1::LUC recordings showed a similarly weak dependence of peak timing on photoperiod, whereas peak timing in the corresponding Col-0 recordings varied more substantially. Although the overall light–dark waveforms of Col-0 and *prr7-11* roots were previously described as similar [43], our analysis focuses on the narrower feature of peak timing relative to dark offset. Together, this model-data agreement suggests that an iPRC inferred solely from 2-h dark-pulse data can capture a *prr7*-associated pattern of peak timing across photoperiods with substantially longer dark intervals. Because the published recordings were obtained in a separate, root-specific experiment and were available only for *prr7-11* among the three mutant lines considered here, this comparison provides cross-condition support for that photoperiod-level pattern rather than quantitative validation of the full duration-dependent PRCs.

PINN-iPRC could also be extended to behavioral and network-level circadian systems, although such applications would require system-specific approaches to phase reconstruction and model formulation. For behavioral circadian rhythms, where singularity-response experiments are difficult to perform, the latent circadian phase would need to be reliably reconstructed from indirect measurements such as locomotor or wheel-running activity. Analytic wavelet analysis [55] and model-based data-assimilation methods [56] offer potential routes to obtaining time-resolved phase estimates, although their reliability during strong resetting requires independent validation. SCN slice recordings present a different modeling challenge: although continuous reporter signals are available, the observed population rhythm emerges from heterogeneous cellular oscillators coupled through the SCN network. Applying PINN-iPRC to such recordings would therefore require a coupled or population-level phase model that accounts for cellular heterogeneity and intercellular coupling.

A more direct application of PINN-iPRC is schedule design in circadian pharmacology [57–59]. PRC-based systems pharmacology studies of the CK1δ/ε inhibitor PF-670462 have shown that circadian drug responses depend on dosing time, environmental light, photosensitivity, and PER2 abundance [60,61]. PINN-iPRC could complement these approaches by inferring how drug exposure modifies instantaneous phase velocity from experimentally accessible cellular reporter trajectories. Once inferred and validated across the relevant dose and exposure conditions, a drug-induced iPRC could be combined with a phase model and a calibrated dose-to-input relationship to compare dosing times, doses, exposure durations, or combined light–drug schedules before more extensive animal or human studies. Such predictions would translate cellular stimulus-response recordings into testable hypotheses for chronotherapeutic schedule design [62].

Several limitations should be understood as identifiability conditions for this physics-informed inverse problem. Because the phase equation constrains the inferred iPRC through the extracted transient phase trajectory, PINN-iPRC depends on reliable phase reconstruction from observed oscillatory signals. Although PINN-iPRC showed relatively stable reconstruction across the tested sample sizes, noise levels, and phase-coverage conditions (Supplementary Fig. S7), weak rhythmicity, strong damping, sparse sampling, measurement noise, or stimulus-induced waveform distortion can still propagate into the learned phase-sensitivity function. This issue was also visible in the synthetic benchmark (Fig. 4), where rapid stimulus-induced phase changes made phase tracking from reporter-like signals imperfect, leading to residual errors in the inferred iPRC. Identifiability also depends on the input term in the phase equation: if the stimulus profile is unknown or poorly approximated, errors in the input can be misattributed to the iPRC. As in other PINN formulations with multiple objective terms, the inferred iPRC may also depend on the balance among data fitting, physics residuals, and regularization terms; inappropriate loss weighting may either over-smooth genuine phase-sensitivity structure or allow noise-driven features to persist [63,64].

The finite-duration predictions in Fig. 6 extend the iPRC inferred from 2-h dark-pulse data to 8-, 12-, and 16-h dark intervals within the same phase-reduction framework. Because the classical iPRC is defined as a local response to weak perturbations near the limit cycle, a phase-sensitivity function inferred from finite or strongly resetting pulses is best interpreted as an effective function within the assumed phase model [48,53]. Importantly, the model does not assume that the final phase shift scales linearly with stimulus duration. The phase shift accumulated over a dark pulse of duration *D* is given by 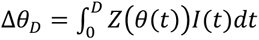, where *I*(*t*) = 1 during darkness and *I*(*t*) = 0 otherwise. Because the phase evolves during stimulation, a longer pulse samples different regions of the iPRC and can therefore alter both the magnitude and shape of the predicted PRC. Thus, the longer-duration PRCs in Fig. 6 are not simple multiples of the 2-h PRC. This qualitative non-proportionality is consistent with human circadian light-response studies in which longer light exposures did not produce proportionally larger phase shifts [65,66].

These extrapolations nevertheless rely on two related assumptions. First, the same phase-sensitivity function is assumed to remain applicable throughout a prolonged stimulus. Substantial amplitude dynamics or other state-dependent departures from a phase-only description may violate this assumption [67]. Second, the effective input is assumed to represent the stimulus consistently across the conditions being compared. At a fixed phase, the first-order phase equation is linear in this effective input: doubling *I*(*t*) doubles the instantaneous perturbation *Z*(*θ*)*I*(*t*). Nevertheless, the accumulated phase shift need not scale linearly with effective input amplitude because the phase trajectory itself changes during stimulation. Here, *I*(*t*) represents the input reaching the oscillator after sensory and signaling transduction rather than the physical stimulus itself. If *S*(*t*) denotes physical stimulus strength, the effective input may be written more generally as *I*(*t*) = ℱ[S](t), where ℱ represents a transformation that may be nonlinear, saturating, or dependent on stimulus history. Consequently, doubling *S*(*t*) need not double *I*(*t*), and maintaining the same physical stimulus does not necessarily imply a constant effective input during prolonged exposure because of adaptation, desensitization, or other time-dependent changes in stimulus transduction. The Arabidopsis data used for inference contain only a single illumination level and a 2-h binary dark-pulse condition and therefore do not determine how the effective input varies across physical stimulus strengths or during prolonged exposure. Accordingly, the calculations in Fig. 6 assume the same illumination intensity and complete-darkness condition as in the inference experiments, with *I*(*t*) = 1 throughout each dark interval and *I*(*t*) = 0 during light. They therefore examine the consequences of changing stimulus duration and temporal schedule rather than physical stimulus intensity or history-dependent transduction.

A natural next step is a prospective, matched-condition evaluation of the predictions in Fig. 6 using the same mutant lines, tissues, reporters, and light intensity as in the iPRC-inference experiments. Single 8-, 12-, and 16-h dark pulses applied at multiple circadian phases would directly test the predicted duration-dependent PRCs. Separate experiments exposing the same lines to repeated 8L:16D, 12L:12D, and 16L:8D schedules would test the predicted peak timing across photoperiods, including the currently untested predictions for *prr5* and *prr9*. Applying each schedule for sufficient cycles from multiple initial phases would further determine whether the trajectories converge to a stable phase relationship, thereby distinguishing phase locking from similarity in observed peak timing. Parallel experiments using calibrated light-to-dark intensity contrasts would help characterize the physical-to-effective input transformation ℱ and determine the range over which first-order input scaling remains valid.

Further methodological advances should improve the reliability of both the extracted phase trajectories and the inferred iPRCs. Abrupt stimulus-induced transitions can introduce phase-extraction errors that propagate directly into PINN-iPRC inference (Fig. 4). Phase-extraction methods that are robust to such transitions, or formulations that jointly infer the phase trajectory and the iPRC from the raw recordings, could reduce this source of error. Future implementations should also incorporate uncertainty quantification that propagates the effects of observation noise, preprocessing choices, phase coverage, regularization strength, and loss weighting to both the inferred iPRCs and predictions under held-out conditions [68–70]. Adaptive or validation-based strategies for balancing data fitting, phase-equation residuals, and regularization could further reduce dependence on manually selected loss weights, as demonstrated in other biological inverse problems [71,72]. Together, these advances would help distinguish robust phase-sensitivity features from inference artifacts and guide experimental design toward stimulus phases, durations, or input profiles that most improve iPRC identifiability.

In summary, PINN-iPRC formulates iPRC inference as a physics-informed inverse problem for extracting hidden phase-sensitivity functions from finite-pulse biological recordings. By combining transient phase trajectories with the phase-equation constraint, PINN-iPRC enables flexible iPRC inference without requiring a full mechanistic state-space model, long continuously perturbed recordings, or specialized singularity-response measurements. More broadly, this study illustrates how PINN-based modeling can use reduced-order dynamical constraints to infer interpretable biological response functions from partial and experimentally accessible observations. The resulting iPRCs provide a basis for testable model-based predictions across perturbation schedules.

## Methods

### **A.** Phase-oscillator formulation and deduced PRC

We modeled the circadian rhythm as a phase oscillator. In the absence of stimulation, the phase *θ*(*t*) ∈ ℝ evolves with a constant angular velocity, 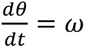, where 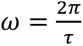 and *τ* is the free-running period.

This unperturbed trajectory was used as the reference for defining stimulus-induced phase shifts.

During stimulation, the phase velocity was assumed to be modified by an external input through a phase-dependent sensitivity function [32,49],

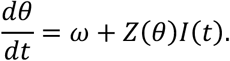

Here, *I*(*t*) denotes the known external stimulus profile and *Z*(*θ*) denotes the infinitesimal phase response curve (iPRC). For pulse perturbations, *I*(*t*) was nonzero only during the stimulation interval.

For an initial phase *θ*_0_ at stimulus onset, we computed the phase shift by integrating the perturbed phase equation over the stimulus duration *T*_stim_. Let *θ*_pert_(*T*_stim_; *θ*_0_) be the perturbed phase at the end of the pulse. The corresponding unperturbed phase is *θ*_0_ + *ωT*_stim_. The phase shift was therefore defined as

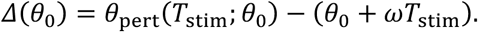

When this phase-shift curve was computed from an inferred iPRC *Ẑ*, we refer to it as the deduced PRC. For comparison with experimentally measured phase shifts, *Δ*(*θ*_0_) was wrapped to the interval [−*π*, *π*), unless otherwise stated.

### **B.** Data

We analyzed two classes of time-series data: synthetic benchmark datasets and experimental plant circadian recordings. Synthetic datasets were used to validate PINN-iPRC against known ground-truth iPRCs, whereas experimental datasets were used to evaluate whether PINN-iPRC can infer phase-response properties from real circadian recordings.

### **B.1.** Synthetic benchmark data generation

Synthetic benchmark datasets were generated by integrating the phase equation

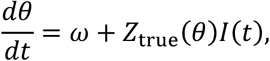

where *Z*_true_ was analytically specified. The stimulus input was modeled as a finite pulse, *I*(*t*) = **1**_[*t*on, *t*on+*T*stim_](*t*), where *t*_on_ is the stimulus onset time, and *T*_stim_ is the stimulus duration. Initial phases at stimulus onset were sampled across [0,2*π*) to cover the full circadian cycle.

The synthetic benchmarks were designed to evaluate iPRC recovery across response regimes of increasing difficulty, including sinusoidal, multi-peaked, and extreme phase-advance cases. In the extreme phase-advance benchmarks, the stimulus was chosen so that the unwrapped transient phase response could approach or exceed one full cycle.

To generate reporter-like observations from the latent phase trajectory, we used

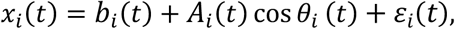

where *x_i_*(*t*) is the observed signal of trajectory *i*, *b_i_*(*t*) is a slowly varying baseline and *A_i_*(*t*) is a slowly varying amplitude envelope, and *ε_i_*(*t*) is zero-mean Gaussian measurement noise with standard deviation *σ* (*σ* = 0 unless otherwise specified; see Supplementary Note 2 for details). This construction allowed us to evaluate the full signal-to-iPRC pipeline, including the effect of phase reconstruction from an observable reporter signal. Thus, unless otherwise stated, PINN-iPRC was trained on phases extracted from *x_i_*(*t*), not on the true latent phase *θ_i_*(*t*).

### **B.2.** Experimental plant circadian time-series

Experimental plant circadian recordings were obtained from the bioluminescence datasets [39]. These datasets measured the responses of Arabidopsis thaliana circadian reporter lines to pulse perturbations. The recordings included CCA1::LUC and TOC1::LUC reporter plants, as well as CCA1::LUC plants carrying *prr5*, *prr7*, or *prr9* mutations. Following the original experimental protocol, plants were entrained under 12 h light/12 h dark cycles before bioluminescence monitoring, and luciferin was supplied before the start of recording. Bioluminescence was then monitored under constant light conditions at 20-min sampling intervals.

For the real-data analyses in this study, we primarily used conventional PRC measurement datasets from *prr5*, *prr7*, and *prr9* CCA1::LUC plants exposed to a 2-h dark pulse. In these experiments, a single 2-h dark pulse was applied once at 72, 78, 84, or 90 h after the start of monitoring, and bioluminescence recording was continued for 7 days. These datasets provided stimulus-response trajectories across different circadian phases and were used as the input data for PINN-iPRC inference. Throughout the Arabidopsis analyses, constant light was treated as the unperturbed condition, with I(t) = 0, and darkness was encoded as I(t) = 1.

Singularity-response datasets from the same study were used only as reference datasets for comparing the SR-based reconstruction. In the original SR protocol, plants were allowed to approach a desynchronized, low-amplitude population state under constant light before receiving a single stimulus at a late recording time. Conventional PRC measurement trajectories and SR measurements were therefore treated as distinct experimental inputs: PINN-iPRC was trained on the conventional stimulus-response trajectories, whereas the SR-based reconstruction used SR information according to the original method [39,40] (see Supplementary Note 1 for details).

Before PINN-iPRC training, each raw bioluminescence trace was converted into a phase trajectory through the preprocessing procedure described below. Each individual plant trace was treated as an independent oscillator trajectory and aligned relative to stimulus onset. Traces with missing values, unstable rhythmicity, or failed phase extraction were excluded during preprocessing.

### **B.3.** Independent photoperiod recordings

Published root CCA1::LUC recordings from *prr7-11* and Columbia-0 (Col-0) plants under 8L:16D, 12L:12D, and 16L:8D conditions [43] were used for an independent comparison of photoperiod-dependent peak timing. The population-mean *prr7-11* traces shown in Fig. 6(e) were digitized from the published recordings, and the corresponding Col-0 recordings provided a wild-type comparison. These data were not used for PINN-iPRC inference. Their analysis and comparison with model predictions are described in Supplementary Note 14.

### **C.** Signal preprocessing and phase extraction

Each raw trace was first interpolated onto a denser time grid to improve the temporal resolution of phase extraction. The interpolated signal was then detrended using Savitzky–Golay filtering [73]. Before phase extraction, the first and last 24 h of each detrended record were removed. Phase trajectories were then obtained from the detrended signals using the Hilbert transform [74], and the resulting wrapped phase was unwrapped over time to obtain a continuous phase trajectory for each individual trace (see Supplementary Note 3 for details).

For perturbation-response inference, the extracted phase trajectories were aligned to stimulus onset. The stimulus onset was set to time zero, and each phase trajectory was shifted onto a common perturbation-aligned time axis. This alignment allowed trajectories stimulated at different absolute recording times to be analyzed together while preserving the phase at which each trajectory received the stimulus. The resulting stimulus-aligned phase trajectories were used as the observed phase input for PINN-iPRC training.

As the phase was extracted from scalar reporter-like signals rather than observed directly, the resulting phase trajectories were treated as estimated phase observations. In particular, finite-pulse stimulation can induce rapid phase progression, amplitude modulation, or transient waveform distortion, which can introduce errors in the reconstructed phase. These residual phase-reconstruction errors were therefore allowed to propagate into the PINN-iPRC input and were considered when interpreting discrepancies between inferred and ground-truth iPRCs in the synthetic benchmarks.

### **D.** Estimation of experimentally measured phase shifts from conventional PRC trajectories

Experimentally measured phase shifts were estimated from conventional PRC measurement trajectories. For each trajectory, peaks or troughs were detected from preprocessed and normalized reporter traces using a slope-change rule. The pre-stimulus rhythm was used to estimate the stimulus-onset phase, and the deviation of post-stimulus peak or trough timing from the expected free-running rhythm was used to estimate the corresponding phase shift (see Supplementary Note 4 for details).

The resulting PRC points were used as experimentally measured phase-shift observations. They served as reference observations for evaluating whether deduced PRCs computed from inferred iPRCs reproduced the observed phase shifts. They were also used in the PRC loss during PINN-iPRC training, where the deduced PRC was constrained to match the observed PRC data at the corresponding stimulus-onset phases.

### **E.** PINN-iPRC model and training objective

PINN-iPRC learns two coupled functions from the extracted phase trajectories [46]. The first network represents the phase trajectory of each observed signal as a function of time, *θ̂_i_* (*t*) = *f_φ_*(*t*)*_i_*, where *i* indexes individual trajectories and *φ* denotes the trainable parameters of the phase-trajectory network. The phase-trajectory network was a fully connected multilayer perceptron with four hidden layers containing 32, 64, 64, and 32 units, respectively, and *N_tra_*_j_ output units. The second network represents the iPRC as a function of phase, *Ẑ*(*θ*) = *g*_=_(sin *θ*, cos *θ*), where *ψ* denotes the trainable parameters of the iPRC network. The sine-cosine input encoding was used to impose periodicity with respect to phase. The iPRC network was a fully connected multilayer perceptron with three hidden layers containing 32 units each and a single output unit. Exponential linear unit (ELU) activations followed every hidden layer, and both output layers were linear.

The model was trained by fitting the extracted phase trajectories, matching the observed finite-pulse phase shifts, and constraining the learned dynamics with the phase equation. The total objective was

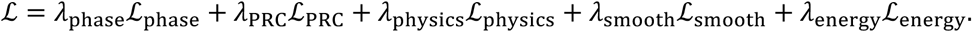

Here, *i* = 1, …, *N*_traj_ denotes individual trajectories, and *j* = 1, …, *T_i_* denotes sampled time points in trajectory *i*. The index *k* = 1, …, *K* labels uniformly spaced phase-grid points on [0,2*π*), and *m* = 1, …, *M* labels experimentally measured phase shifts.

The phase data loss fitted the neural phase trajectories to the extracted unwrapped phases,

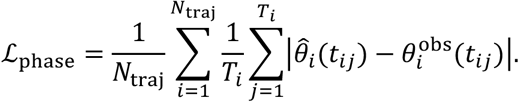

For PRC alignment, experimentally measured phase-shift observations 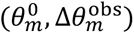 were used as targets; the learned iPRC was evaluated at the same initial phases 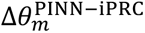, and the resulting deduced phase shifts 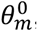 were compared with the observed values,

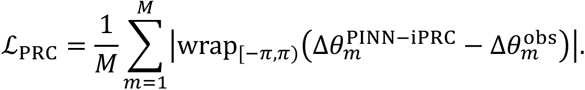

The physics loss penalized deviations from the phase equation,

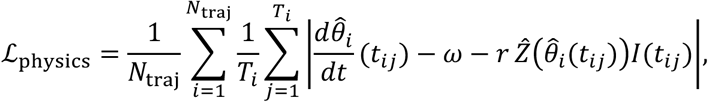

where the derivative *dθ*^S^*_i_*/*dt* was computed by automatic differentiation. The calibration factor *r* was shared across all trajectories within each dataset and applied only in the physics residual to accommodate scale differences in the reconstructed transient phase dynamics. It was initialized at 1 and constrained to 0.3 < *r* < 3.0. PRC integration and all reported predictions used the unscaled iPRC-network output *Ẑ*. This term enforces that the learned phase trajectory evolves according to the phase equation under the known stimulus profile, thereby linking the neural iPRC to time-resolved phase dynamics rather than only to endpoint phase shifts. Smoothness of the inferred iPRC was imposed on a uniform phase grid 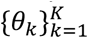 covering [0,2*π*). With *Ẑ_k_* = *Ẑ*(*θ_k_*) and Δ*θ* = 2*π*/*K*,

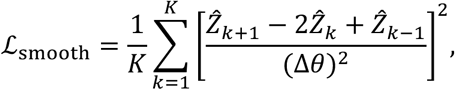

with circular indexing *Ẑ*_0_ = *Ẑ_K_* and *Ẑ_K_*_+1_ = *Ẑ*_1_.

The energy penalty, when enabled, controlled the overall magnitude of the inferred iPRC,

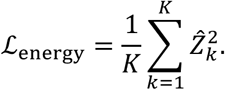

Loss weights were selected empirically in preliminary runs based on stable convergence of the phase-trajectory and physics-residual losses and, where applicable, the PRC-alignment loss. The phase-data weight was kept large to prioritize agreement with the extracted phase trajectories. The smoothness weight was increased when smoother iPRC profiles were required, whereas the energy weight controlled the overall magnitude of the inferred iPRC. The PRC-alignment weight was adjusted according to the reliability and intended strength of marker-based PRC alignment. Neither SR-based iPRC estimates nor known synthetic ground-truth iPRCs were used to select or tune these weights. The selected weights were then held fixed across all independent initializations for each dataset, with no seed-specific retuning. All training settings were recorded with the corresponding model outputs. For the main analyses in Figs. 4 and 5, we used *λ_p_*_ℎ*ase*_ = 80,000, with dataset-specific weights of *λ_p_*_ℎ*ysics*_ = 4,000 − 80,000, *λ_PRC_* = 1,000 − 8,000, *λ_smoot_*_ℎ_ = 1 − 1,000, and *λ_ener_*_g*y*_ = 1 − 100. Detailed network architectures, hyperparameters, loss weights, and optimization settings are provided in Supplementary Note 5 and Supplementary Table S3. We also performed sensitivity analyses by varying the smoothness and energy regularization weights, as described in Supplementary Note 10 (Supplementary Fig. S6).

The phase-trajectory network, iPRC network, and trainable calibration factor *r* were jointly optimized in 32-bit floating-point precision using Adam. All parameter groups used a constant learning rate of 10^–3^ with a weight decay of 10^–3^. The stopping criterion was completion of a prespecified fixed epoch budget; no early stopping or validation-based model selection was used. Training was performed for 5,000 epochs for the sinusoidal synthetic benchmark, 10,000 epochs for the multi-peaked and extreme synthetic benchmarks, and 5,000 epochs for each experimental dataset. No learning-rate scheduling, PRC-loss warm-up, or gradient clipping was used.

PINN optimization used only the interval from 3 h before pulse onset to 3 h after pulse offset, *t* ∈ [−3, *T_stim_* + 3] (hour). This interval defines the PINN fitting window; phase preprocessing and marker-based PRC estimation used the longer surrounding recordings. Time was rescaled as *t* = *t*/*t_max_*, where *t_max_* = *T_stim_* + 3 (hour) in the reported analyses, and automatic-differentiation derivatives were divided by *t_max_* to recover rad ℎ^–1^. After fourfold interpolation, the training interval contained 97 time points per trajectory for the 2-h-pulse analyses and 145 points per trajectory for the 6-h case. Full-batch optimization was used, with losses accumulated over all trajectories before one optimizer update per epoch. No additional randomly sampled collocation points or validation split was used.

Each benchmark and mutant analysis shown in Figs. 4 and 5 was repeated using 20 independent random initializations. The reported central iPRC was the pointwise arithmetic mean of the final raw iPRC-network outputs *Ẑ*(*θ*), and the corresponding PRC was calculated by integrating this ensemble-mean iPRC. Across seeds, the phase-averaged pointwise sample standard deviation was 1.0–1.6% of the peak-to-peak ensemble-mean iPRC amplitude for the synthetic benchmarks and 0.54%, 1.28%, and 8.58% for *prr5, prr9*, and *prr7*, respectively. Fig. 6 reused the ensemble-mean iPRCs inferred for Fig. 5 and therefore involved no additional stochastic fitting. Case-specific trajectory numbers, epoch budgets, ensemble sizes, and loss weights are reported in Supplementary Table S3, with further implementation details provided in Supplementary Note 5.

## Data Availability

Raw experimental data used in this study are derived from previously published datasets [39,43]. The processed datasets required to reproduce the main figures are provided with the submission as part of the accompanying code package and will be made publicly available upon publication.

## Code Availability

The source code required to reproduce the main figures is provided with the submission and will be made publicly available upon publication.

## Acknowledgements

The research was supported by the following agencies and institutions: the Institute for Basic Science (grant no. IBS-R029-C3, J.K.K.); Samsung Science and Technology Foundation (grant no. SRFC-IT2601-01, J.K.K.); the National Research Foundation of Korea (NRF) grant funded by the Korean government (MSIT) in 2026 (grant no. RS-2026-25613293, J.K.K.).

## Author Contributions Statement

Simulation: D.W.

Analysis of results: All authors First draft: All authors Writing: All authors

